# Suppression of Pectinase Genes Confers Stable Enhancement of Fruit Firmness through Modulation of Pectin Structure in Strawberry

**DOI:** 10.64898/2026.01.29.699421

**Authors:** Pablo Ric-Varas, José A. Mercado-Hornos, Julia Schückel, Sara Aguado, Marta Barceló, J. Paul Knox, Rosario Blanco-Portales, Juan Muñoz-Blanco, Antonio J. Matas, Candelas Paniagua, Miguel A. Quesada, Sara Posé, José A. Mercado

**Affiliations:** Instituto de Hortofruticultura Subtropical y Mediterránea ‘La Mayora’ (IHSM-UMA-CSIC), Departamento de Botánica y Fisiología Vegetal, Universidad de Málaga, 29071, Málaga, Spain; Department of Plant and Environmental Sciences, University of Copenhagen, Copenhagen, Denmark; IFAPA Centro de Málaga, Cortijo de la Cruz s/n, 29140, Málaga, Spain; Centre for Plant Sciences, Faculty of Biological Sciences, University of Leeds, Leeds, LS2 9JT, UK; Departamento de Bioquímica y Biología Molecular, Universidad de Córdoba, 14071, Córdoba, Spain

**Author notes:** These authors contributed equally to this work and share first authorship (Pablo Ric-Varas, José A. Mercado-Hornos).

**Keywords:** *Fragaria*, cell wall, fruit softening, fruit ripening, fruit texture, pectins

## Abstract

Fruit softening is primarily determined by modifications in the cell wall architecture, which are mediated by the coordinated activity of cell wall-degrading enzymes. In strawberry (*Fragaria × ananassa* Duch.), transgenic suppression of the genes encoding the pectinases polygalacturonase (PG), β-galactosidase (βGal), and rhamnogalacturonan lyase (RGLyase), reduced fruit softening. In this study, we evaluated the fruit firmness phenotype of selected transgenic lines across several harvest years, from 3 to 9 years depending on the line, and analyzed the cell wall composition of ripe fruits using a carbohydrate microarray. Multiple linear regression analysis revealed that fruit firmness was significantly affected by both genotype and harvest year, while fruit size and soluble solids content showed no significant contribution. All pectinase-silenced lines exhibited increased firmness relative to the wild type, with those lines with PG down-regulated showing the most significant effects, followed by B-Gal and RGLyase fruits. The firmer phenotype was maintained stably in all the transgenic lines during the different years analyzed. Carbohydrate microarray analyses of sequentially extracted cell wall fractions demonstrated that transgenic ripe fruits retained higher levels of low methyl-esterified homogalacturonan (HG) and rhamnogalacturonan I (RG-I) epitopes compared to wild-type ripe fruits, resembling the composition of white-stage control fruits. Principal component analysis of microarray data revealed a clear separation between wild-type ripe fruits and transgenic lines, with the latter clustering near the earlier developmental stages of wild-type fruits. Correlation analysis further revealed positive associations between increased firmness and the abundance of high-methylated HG pectic epitopes in the water fraction, recognized with JIM7, and low-methylated HG abundance in the rest of the fractions (JIM5, LM18, and LM19). Overall, these results suggest that suppressing pectinase genes alters pectin remodeling during ripening, resulting in the retention of structurally intact pectin domains and increased fruit firmness. These genes are therefore excellent candidates for the improvement of this key quality trait in strawberry.

## 1. Introduction

The cultivated strawberry (*Fragaria* × *ananassa* Duch.) is one of the most economically important fruit crops worldwide, with a global production of 10.4 million tonnes harvested from 435,000 hectares in 2023 (FAOSTAT, 2025). In addition to their attractive sensory attributes, strawberries are a rich source of bioactive and antioxidant compounds, contributing to their nutritional and health-promoting value (Newerli-Guz et al., 2023; Charoenwoodhipong et al., 2024). Furthermore, *F. × ananassa* and its diploid progenitor *F. vesca* are widely used as model systems for investigating the ripening process in non-climacteric fruits (Vondracek et al., 2024). Strawberries belong to the group of soft fruits, which also includes raspberries and blackcurrants (Manning, 1993). These fruits undergo rapid softening during ripening, resulting in a melting texture that severely limits their postharvest handling, long-distance transport, and shelf life, leading to substantial economic losses. Fruit softening is associated with the disassembly of the middle lamella and primary cell wall, which reduces wall strength and intercellular adhesion, and is accompanied by a decrease in cell turgor (Brummell et al., 2022). In strawberries, cell wall modification is the primary determinant of softening (Wang et al., 2018), whereas in other fruits such as grapes and tomatoes, postharvest water loss plays a more prominent role (Saladié et al., 2007; Wada et al., 2009).

Cell wall remodeling during fleshy fruit softening involves depolymerization of pectins, particularly homogalacturonan (HG), and xyloglucans, demethylesterification of HG, and the loss of galactan and arabinan side chains from rhamnogalacturonan I (RG-I) (Brummell et al., 2022). In strawberry, there is an extensive solubilization of pectins during ripening, with a marked increase in water- and chelator-extractable fractions and a concomitant decrease in tightly bound fractions (Paniagua et al., 2017a). This process is driven by pectin depolymerization mediated by polygalacturonases (PGs), pectate lyases (PL), and other pectinases; the release of neutral sugars from RG-I, which increases cell wall porosity and enzyme accessibility; and progressive demethylesterification of HG by pectin methylesterases (PMEs), leading to pectin network loosening through electrostatic repulsion (Brummell, 2006; Posé et al., 2019).

Genetic manipulation of pectinase-encoding genes has provided direct evidence for their contribution to fruit softening (Brummell et al., 2022; Shi et al., 2023). Silencing of genes acting on HG (PG, PL, PME) or RG-I (rhamnogalacturonan lyase, β-galactosidase) has been performed in tomato, strawberry, and apple, often resulting in reduced softening and extended shelf life. However, these studies typically focus on single genes, and differences in genetic backgrounds, growth conditions, and analytical methods make it difficult to compare the specific contributions of individual enzymes to cell wall disassembly and fruit softening.

The strawberry cultivar ‘Chandler’, released by the University of California, Davis in 1983 (Chandler et al., 2012), produces fruits with outstanding organoleptic quality but limited postharvest life due to soft texture. Although it was largely replaced in the 1990s by firmer cultivars, ‘Chandler’ remains profitable for processed products (frozen fruits, juices) and as a genetic background for functional studies. Over the past 15 years, a collection of transgenic ‘Chandler’ lines with silenced pectinase genes (e.g., PG, PL, β-Gal, RGLyase) has been generated, exhibiting varying degrees of increase in firmness compared to wild-type fruits (Jiménez-Bermúdez et al., 2002; Paniagua et al., 2016, 2020; Ric-Varas et al., 2024). The present study aimed to conduct a meta-analysis of fruit quality variables across these transgenic lines over multiple growing seasons to assess the impact of specific pectinases on fruit softening. In addition, the glycome profiling using carbohydrate microarrays previously performed in some of these cell wall mutants (Paniagua et al., 2020; Ric-Varas et al., 2024) was reanalysed to identify potential glycan fingerprints associated with strawberry softening.

## 2. Materials and methods

### 2.1. Plant Material

Strawberry plants, *Fragaria* × *ananassa* Duch. cv. ‘Chandler’, were used. The transgenic strawberry genotypes were obtained and described in previous research papers and are listed in Table 1. These lines were obtained through the *Agrobacterium tumefaciens*-mediated transformation of leaves from in vitro strawberry plants following the protocol described by Barceló et al. (1998). Antisense *FaPG1* lines (PGI) were generated in 2005; antisense *FaPG2* (PGII) and stacking of antisense *FaPG1* and *FaPG2* (PGI/II) in 2011; antisense *Fa*β*Gal4* (β-Gal) in 2010, and, finally, *FaRGLyase1* RNAi lines (RG) in 2014. All transgenic lines exhibited a downregulation of the corresponding target gene of greater than 90% in red-ripe fruit compared to control, non-transgenic fruits, as assessed by qRT-PCR (Paniagua et al., 2016, 2020; Ric-Varas et al., 2024).

**Table 1.** List of transgenic lines used in this research.

| Transgenic genotype | Pectinase activity modified | Gene | Reference |
| --- | --- | --- | --- |
| PGI-29<br>PGI-62 | Polygalacturonase | Antisense of *FaPG1* | Quesada et al. (2009) |
| PGII-5<br>PGII-8 | Polygalacturonase | Antisense of *FaPG2* | Paniagua et al. (2020) |
| PGI/II-9<br>PGI/II-16 | Polygalacturonase | Stacking of antisense sequences of *FaPG1* and *FaPG2* | Paniagua et al. (2020) |
| RG26<br>RG87 | Rhamnogalacturonan lyase | RNAi of *FaRGLyase1* | Ric-Varas et al. (2024) |
| β-Gal28<br>β-Gal37 | β-galactosidase | Antisense of *FaβGal4* | Paniagua et al. (2016) |
| Acel20 | endo-β-(1,4)-glucanase | Antisense of *FaEG3* | Mercado et al. (2010) |

**Table 2.** Significant correlations between increment in fruit firmness and mAb signal.

| Epitope | Cell wall Antibody | Cell wall fraction | Pearson correlation coefficient | *P* |
| --- | --- | --- | --- | --- |
| Low methyl-esterified HG | JIM5 | Water | -0.766 | 0.004 |
|  |  | KOH | 0.673 | 0.016 |
|  | LM18 | Sodium carbonate | 0.610 | 0.035 |
|  |  | Cadoxen | 0.669 | 0.017 |
|  | LM19 | Cadoxen | 0.638 | 0.025 |
| Partially methyl-esterified HG | JIM7 | Water | 0.703 | 0.018 |
| (1→4)-β-D-galactan, RG-I | LM5 | Cadoxen | 0.587 | 0.044 |
| XXXG/galactosylated xyloglucan | LM25 | Cadoxen | -0.649 | 0.025 |

The pectinase transgenic lines analyzed in this study were evaluated phenotypically over a period of three to nine harvest years depending on the line (Supplementary Table 1), from 2011 to 2024. Control and transgenic plants were propagated by runners each year, and only fruits produced by one-year-old plants were evaluated. Plants were grown in 22 cm pots containing a mixture of peat moss, sand, and perlite (6:3:1) and cultured in a confined greenhouse equipped with a cooling system to maintain a maximum temperature below 30 °C. Average monthly temperatures ranged from 15°C in January to 24°C in June. Plants were fertilized weekly with Nitrofoska® solub 12-5-30 (EuroChem Agro Iberia S.L., Barcelona, Spain).

### 2.2. Phenotypic analysis of transgenic fruits

For phenotypic evaluation, control and transgenic fruits were collected at the fully ripe stage, with 100% surface red, from March to June (Supplementary Fig. 1). Fruit weight, length, and width were recorded. A refractometer (Atago N1) was used to measure soluble solids, and a handheld penetrometer (Effegi) with a 9.62 mm² cylindrical needle was used for firmness measurements. The number of fruits assessed each year is displayed in Supplementary Table 1. All transgenic lines were evaluated during at least three different years.

### 2.3. Cell wall extraction

For cell wall analysis, fruits from control plants were harvested at three developmental stages: green-unripe, white-unripe (full white receptacle), and ripe (full red receptacle). Red fruits from the transgenic line Cel20, expressing an antisense sequence of the endo-β-Glucanase gene *FaEG3*, were used as an additional control, since this line did not show any alteration in fruit firmness (Mercado et al., 2010). In the case of pectinase lines, only red-ripe fruits were employed. All fruits were frozen in liquid nitrogen immediately after harvest and stored at −40 °C until use. Samples for cell wall extraction were collected in 2012 and 2015, except for PGI-62 that were collected in 2011, and in all cases during the same period of the year (late May to early June).

Frozen fruits were de-acheneized and finely milled into powder under liquid nitrogen. Ten g of fruits were extracted with 20 ml of PAW (phenol: acetic acid: water, 2:1:1, w/v/v) as described by Redgwell et al. (1992) to prevent cell wall-degrading enzymes from acting during extraction and further fractionation. After centrifugation at 4000 *g*, the pellets were de-starched by treatment with 90% aqueous DMSO, and the final residue, considered the cell wall extract, was lyophilized. Cell wall was sequentially fractionated in different solvents: water, 0.1 M Na_2_CO_3_, 4 M KOH, and cadoxen (31% 1,2-diaminoethane with 0.78 M cadmium oxide (v/v)). The last two solvents included 0.1% NaBH freshly added before use. To this end, 10 mg of cell wall material was homogenized in a tissue lyser with 500 μL of water at 30 Hz for 20 min, followed by gentle stirring for one hour at room temperature. Then, the extracts were centrifuged at 2700 g for 15 min, and the supernatant was stored as the water fraction. The pellet was further extracted with a successive solvent, following the same steps.

### 2.4. Microrray of cell wall fractions

Samples of each fraction were printed simultaneously on the same sheet of nitrocellulose as adjacent arrays using an ArrayJet Sprint (ArrayJet, Roslin, UK) and quantified as previously described (Kračun et al., 2017). Briefly, the printed nitrocellulose sheets were probed with the primary monoclonal antibodies (mAbs) diluted (1/10) in phosphate-buffered saline (PBS) containing 5% (w/v) milk powder (MPBS). Secondary anti-rat or anti-mouse antibodies conjugated to alkaline phosphatase (Sigma) were diluted (1/5000) in MPBS. Supplementary Table 2 shows the primary mAbs used in this study. All of them were obtained from PlantProbes (Leeds, UK) except RU1 and RU2, which were kindly supplied by M.C. Ralet (Biopolymères Interactions Assemblages, Nantes, France). Developed microarrays were scanned (CanoScan 8800F), converted to TIFFs, and signals were processed by ImaGene 6.0 microarray analysis software (BioDiscovery), as described by Moller et al. (2007). The mean spot signals obtained from four experiments are presented in heat maps, in which color intensity is correlated with signal. The highest signal in each dataset was set to 100, and all other values were normalized accordingly. A cut-off value of 5 was applied.

### 2.5. Statistical analysis

To assess the influence of genotype on fruit firmness (response variable), a multiple linear regression model was fitted, including adjustments for several covariates, such as fruit length, width, soluble solids content, and harvest year. Each fruit was considered as an independent biological replicate. For each year, only genotypes with at least 8 independent fruit samples were included in the analysis. Interaction terms were not included in the model. All numerical variables were standardized to have a mean of 0 and a standard deviation of 1 to ensure that the estimated coefficients were comparable across variables. The model’s diagnosis was performed graphically by plotting the residuals against the fitted values to check for constant variance and by examining the Q-Q plot of the residuals to verify normality. The Box-Cox method was employed to investigate potential transformations of the response variable that best fit the data. To contrast the null hypothesis that the estimated coefficients of the model were equal to 0, a *t*-test was used, and P-values were adjusted using the Benjamini–Hochberg method. The F-test was used to compare the model with all predictors with a model that included only a subset of predictors.

In the carbohydrate microarray experiment, normalized data were subjected to principal component analysis (PCA) using R. Only those mAbs detected in at least three different samples per fraction were included in the analysis. All tests were performed at a significance level of P = 0.05.

## 3. Results

### 3.1. Effect of harvest year on firmness of transgenic ripe fruits

Bivariate plots showed no clear relationship between fruit firmness and other measured traits such as fruit weight, length, width, and soluble solids content, with low correlation coefficients (r < 0.09; Supplementary Figure 2). In contrast, fruit firmness varied noticeably among genotypes and harvest years (Figure 1A). To assess the relative effects of these variables on fruit firmness, the data were fitted to a multiple linear regression model. Fruit weight was excluded to avoid collinearity, as it was strongly correlated with fruit length and width (r = 0.79 and r = 0.74, respectively; both P < 0.001; Supplementary Figure 2). The residuals of the fitted model showed deviations from linearity at the extremes of the Q–Q plot (Supplementary Figure 3), and the Box-Cox analysis suggested that a transformation of the response variable could improve model performance (95% CI for λ = 0.417-0.582). Consequently, the model was refitted using the square root of fruit firmness as the response variable, and all numerical predictors were standardized to allow direct comparison of estimated coefficient magnitudes.

**Figure 1.**
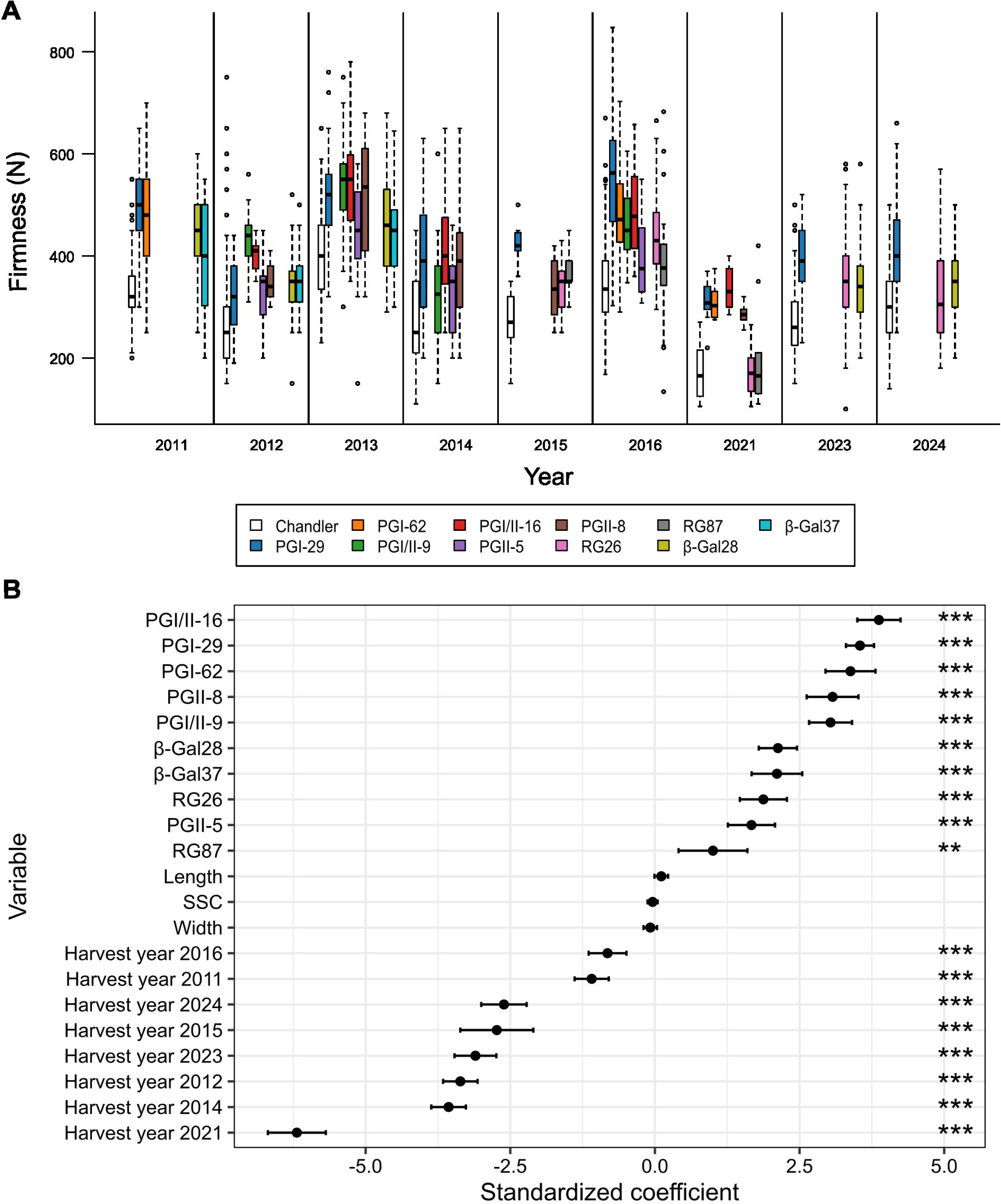
Influence of genotype in fruit firmness across different harvest years. (A) Boxplots showing fruit firmness in ‘Chandler’ and transgenic lines in different harvest years. The line in the box indicates the median; upper and lower bounds indicate the 25th and 75th percentiles; whiskers show 1.5 × interquartile range; and values outside are shown as individual data points. (B) Multiple linear regression model using square root of fruit firmness as response variable and fruit length, width, soluble solid content, harvest year and genotype as response variables. Points represent the estimate coefficient and error bars the 95% confidence interval. For each coefficient, adjusted P-value (Benjamini–Hochberg method) of the t-test is shown. A significant difference was considered if *P* < 0.05.

The fitted model explained nearly half of the variability in fruit firmness (adjusted R² = 0.481), and the assumptions of homoscedasticity and residual normality were met (Supplementary Figure 3). Estimated coefficients for fruit length, width, and soluble solids content were not significant (Figure 1B). Furthermore, a reduced model including only harvest year and genotype did not differ significantly from the complete model (F-test, P = 0.180), indicating that fruit length, width, and soluble solids content did not significantly contribute to explaining fruit firmness in this dataset.

All transgenic genotypes exhibited significantly positive coefficients (Figure 1B), indicating that their fruits were firmer than those of the wild-type (WT, control non-transformed plants) reference level. Among the transgenic lines, PGI/II-16, PGI-29, PGI-62, PGII-8, and PGI/II-9 showed the most significant positive effects on fruit firmness, whereas RG87 and PGII-5 had the most negligible effect. For each harvest year, the increment in fruit firmness relative to the control was calculated and the average values of fruit firmness increments are shown in Fig. 2.

**Figure 2.**
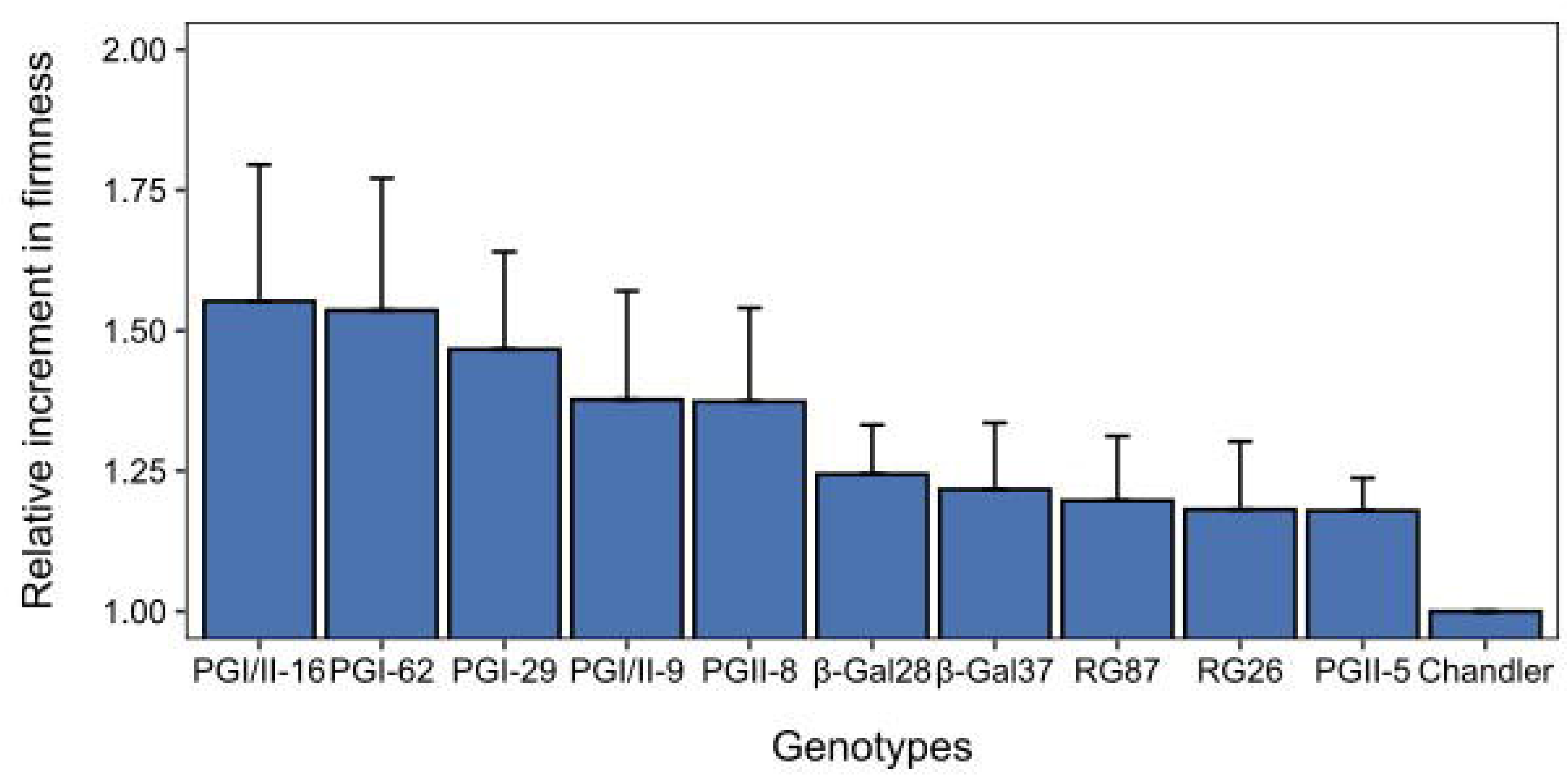
Average increase in firmness in the different transgenic lines evaluated. For each harvest year, the increment in fruit firmness relative to the control line was calculated and the bars correspond to mean±SD of these increment values.

Regarding the harvest year, all years had negative coefficients, suggesting that fruits harvested in 2013 (the reference level) were the firmest. Conversely, fruits from the 2021 harvest had the lowest estimated firmness values.

### 3.2. Cell wall remodeling in transgenic ripe fruits

Cell wall material from the ripe fruits of transgenic plants with low expression levels of pectinase genes was isolated, fractionated, and analyzed using a carbohydrate microarray. Wild type (WT, control non-transgenic) strawberry fruits at different developmental stages, green-unripe, white, and red-ripe, as well as ripe fruits from the antisense endo-β-(1,4)-glucanase gene *FaEG3* were also included. This transgenic line displayed similar fruit firmness to the control (Mercado et al., 2010). The solvents used for extractions yielded fractions enriched in pectins loosely and tightly bound to the cell wall (water and sodium carbonate fractions, respectively), xyloglucans (4 M KOH), and matrix polysaccharides highly imbricated into cellulose microfibrils (cadoxen). The distribution of monoclonal antibodies (mAbs) showing a quantifiable signal in the microarray is summarized in Fig. 3. Seven mAbs were common to all fractions: LM18 and LM19 (low methyl-esterified HG); RU1 and RU2 (RG-I backbone); LM5 and LM6-M (galactan and arabinan RG-I side chains, respectively); and JIM13 (arabinogalactan proteins, AGPs). LM7, which recognizes partially methyl-esterified HG in a non-blockwise pattern, was specific to the sodium carbonate (SC) fraction. LM24 (galactosylated xyloglucans), LM10 (xylans), and LM21 (heteromannans) were detected only in the cadoxen fraction. Finally, JIM7 (methyl-esterified HG) was present only in the water fraction, as subsequent chemical extractions (SC and beyond) de-esterified pectins.

**Figure 3.**
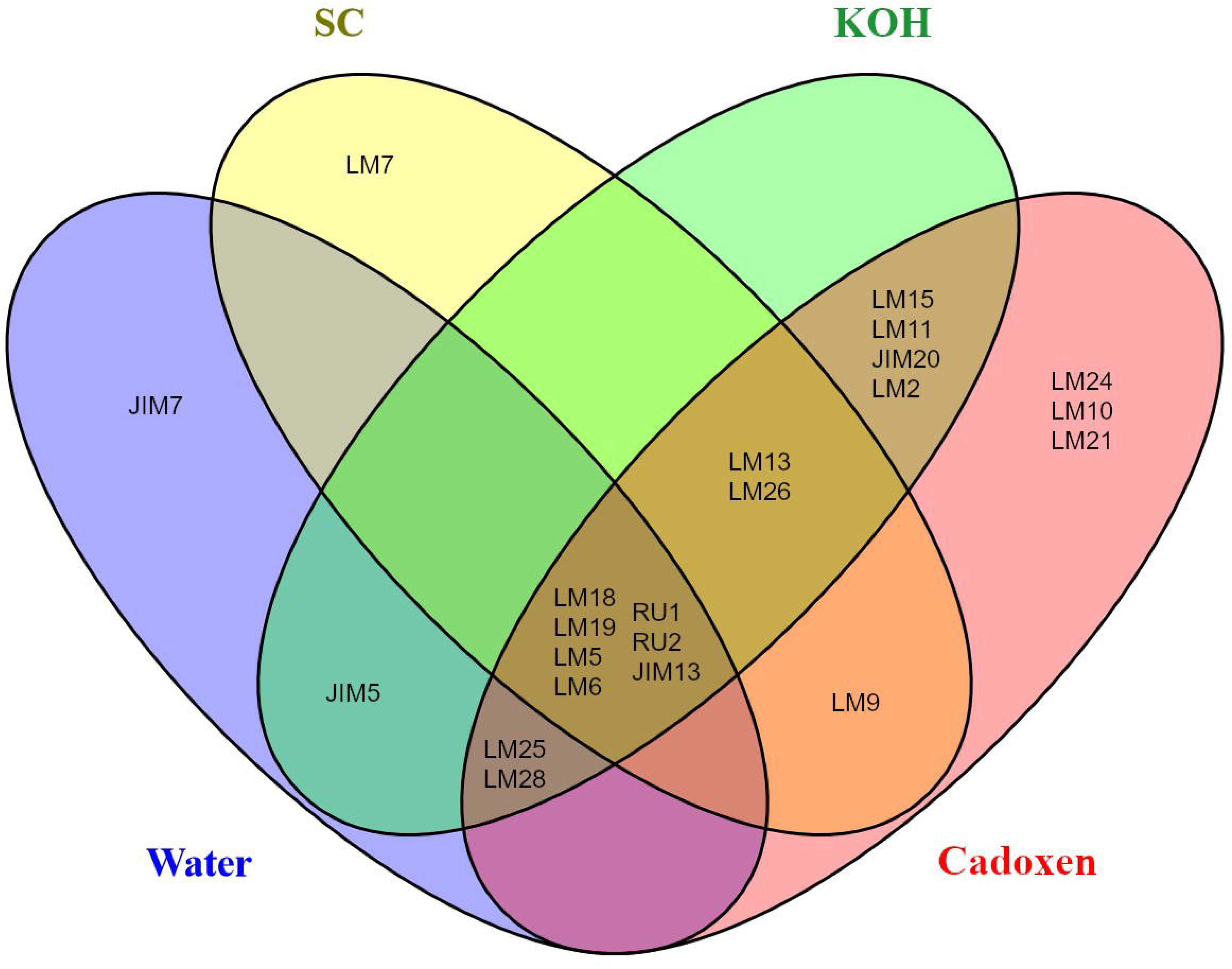
Veen diagram of mAb detected in the different cell wall fractions.

Figure 4 shows the heatmap of the relative abundance of cell wall epitopes in the water and sodium carbonate fractions, while KOH and cadoxen fractions are represented in Fig. 5. In the whole microarray, antisense endo-β-glucanase line Cel20 displayed a mAb signal pattern similar to control ripe fruits. In the water fraction of control fruits, absorbances for low methyl-esterified HG (LM18, LM19, and JIM5) were weak in unripe fruits but increased substantially in red-ripe fruits, particularly for the JIM5-recognized epitopes. Conversely, the JIM7 signal (methyl-esterified HG) peaked at the white stage and declined sharply in ripe fruits. Pectinase lines exhibited JIM7 signals comparable to WT at the white stage, particularly in PG and RGLyase mutants. However, JIM5-recognized demethylated HG epitopes were less abundant in transgenic ripe fruits than in WT controls. RG-I backbone (RU1, RU2) and side chain (LM5, LM6-M) epitopes increased in WT fruits from green to white stages but were undetectable in ripe fruits. In contrast, transgenic ripe fruits retained substantial levels of these epitopes, comparable to those of white-stage control fruits. The strongest RG-I-related signals were observed in the RGLyase mutants.

**Figure 4.**
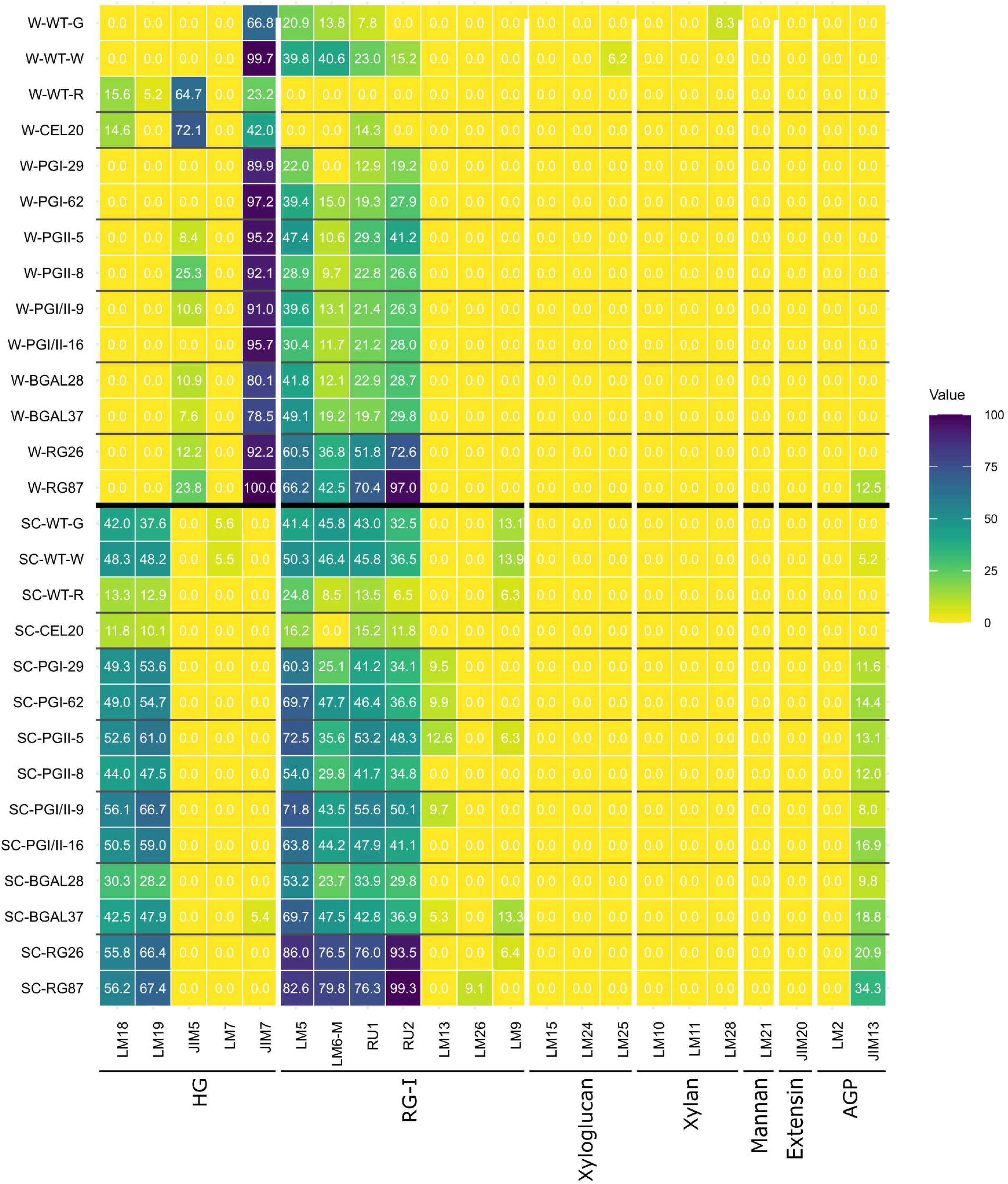
Heat map illustrating the relative abundance of cell wall epitopes detected by the set of mAbs in the water (W) and sodium carbonate (SC) fractions. The highest spot signal was set to 100, and all remaining values were normalized accordingly.

**Figure 5.**
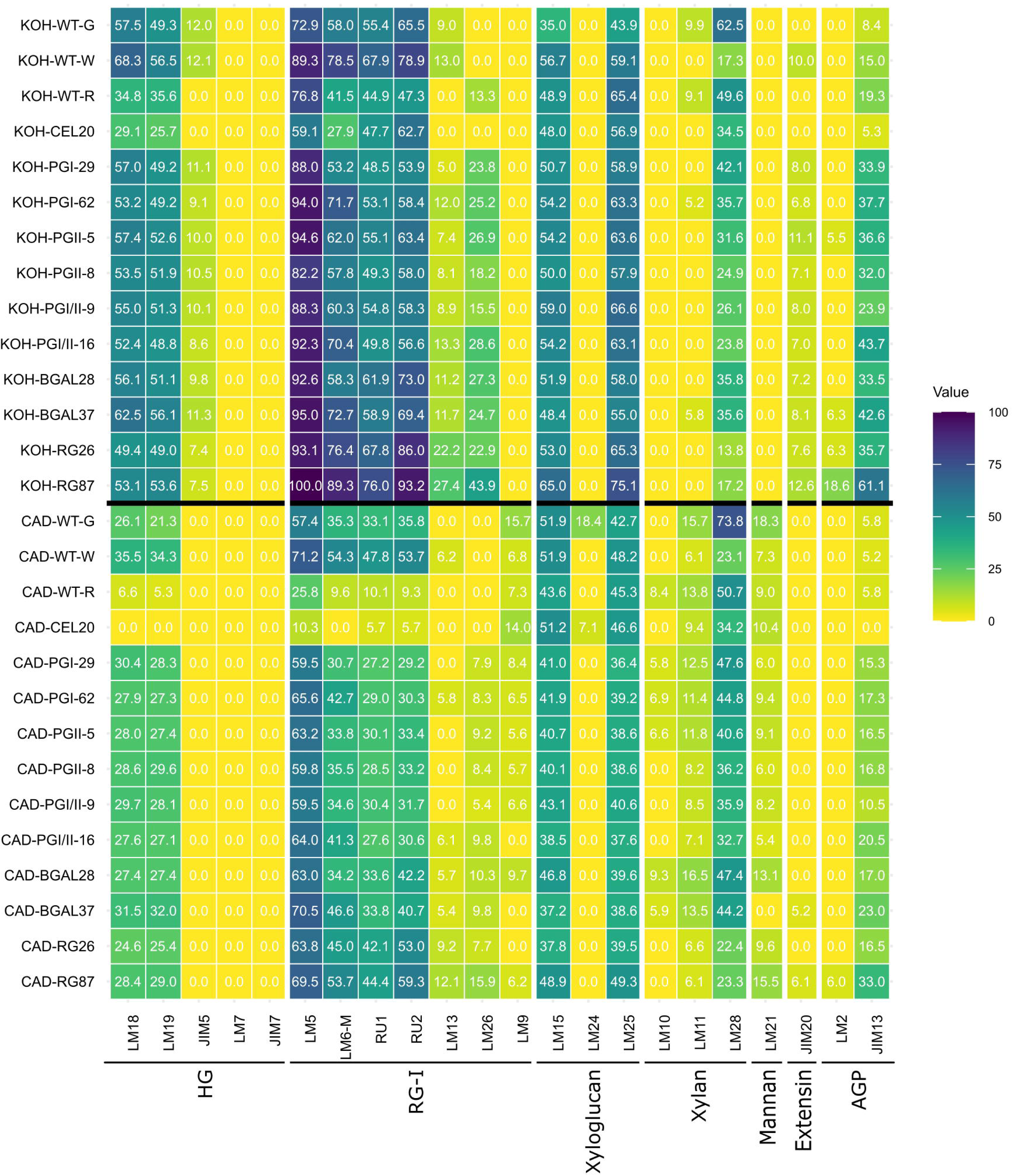
Heat map illustrating the relative abundance of cell wall epitopes detected by the set of mAbs in the KOH and cadoxen (CAD) fractions. The highest spot signal was set to 100, and all remaining values were normalized accordingly.

In the SC fraction, both HG and RG-I epitope signals decreased at the ripe stage compared to the green and white stages, with minor differences observed among pectinase lines and WT fruits at the unripe stages. As in the water fraction, RG-I epitopes were particularly abundant in RGLyase mutant lines.

The KOH and cadoxen fractions showed similar epitope distribution patterns. Both were enriched in xyloglucans, detected at comparable levels by LM15 and LM25. Xyloglucan abundance remained stable across fruit development in WT and between transgenic and WT lines. Both fractions also contained considerable amounts of pectins (HG and RG-I), which peaked at the white stage in WT and declined at ripening. In contrast, transgenic lines maintained pectin levels comparable to those of unripe fruits. Xylans, mainly detected by LM28, were most abundant at the green stage in WT. Transgenic lines generally showed xylan levels similar to WT ripe fruits, except for RGLyase mutants, which displayed lower levels. Extensin epitopes (JIM20) were detected in the KOH fraction, whereas mannans (LM21) were localized in the cadoxen fraction, with both polymer types showing low signal intensity. All transgenic lines exhibited higher AGP mAb signals (particularly JIM13) than WT, most prominently in the KOH fraction.

Microarray data were subjected to principal component analysis (PCA). The first two principal components explained a high proportion of total variance, from 91% in the case of water to 67.4% in cadoxen. In the water fraction, all transgenic lines with pectinase genes down-regulated clustered near WT fruits at green and white stages, except for RGLyase lines, which were displaced along the positive axis of PC1 (Fig. 6A). This dimension was positively associated with RG-I mAbs and negatively with methyl-esterified HG (JIM7). Ripe control fruits were clearly separated from other samples due to their high JIM5 (low methyl-esterified HG) signal. A similar pattern was observed in the SC fraction. In this case, green and white fruits were separated from transgenic PG and B-Gal lines along PC2, mainly influenced by LM9 (feruloylated galactan) and LM13 (AGP glycan) epitopes (Fig. 6B). PCA plots for KOH and cadoxen fractions were also comparable (Fig. 7). In control fruits, developmental stages were separated clearly along both PCs, reflecting compositional changes in these fractions during ripening. Pectinase-silenced lines were positioned closer to white-stage control fruits than to green or ripe stages. Among them, RG87 displayed the most distinct KOH and cadoxen profiles, shifted toward positive PC1 scores due to higher RG-I epitope content. As expected, the endo-β-glucanase Cel20 line grouped close to the WT at the red stage in the PCA analysis, except for cadoxen fraction, where it was displaced to the negative PC2 axis mainly due to its higher signal of RG-I epitope recognized by LM9.

**Figure 6.**
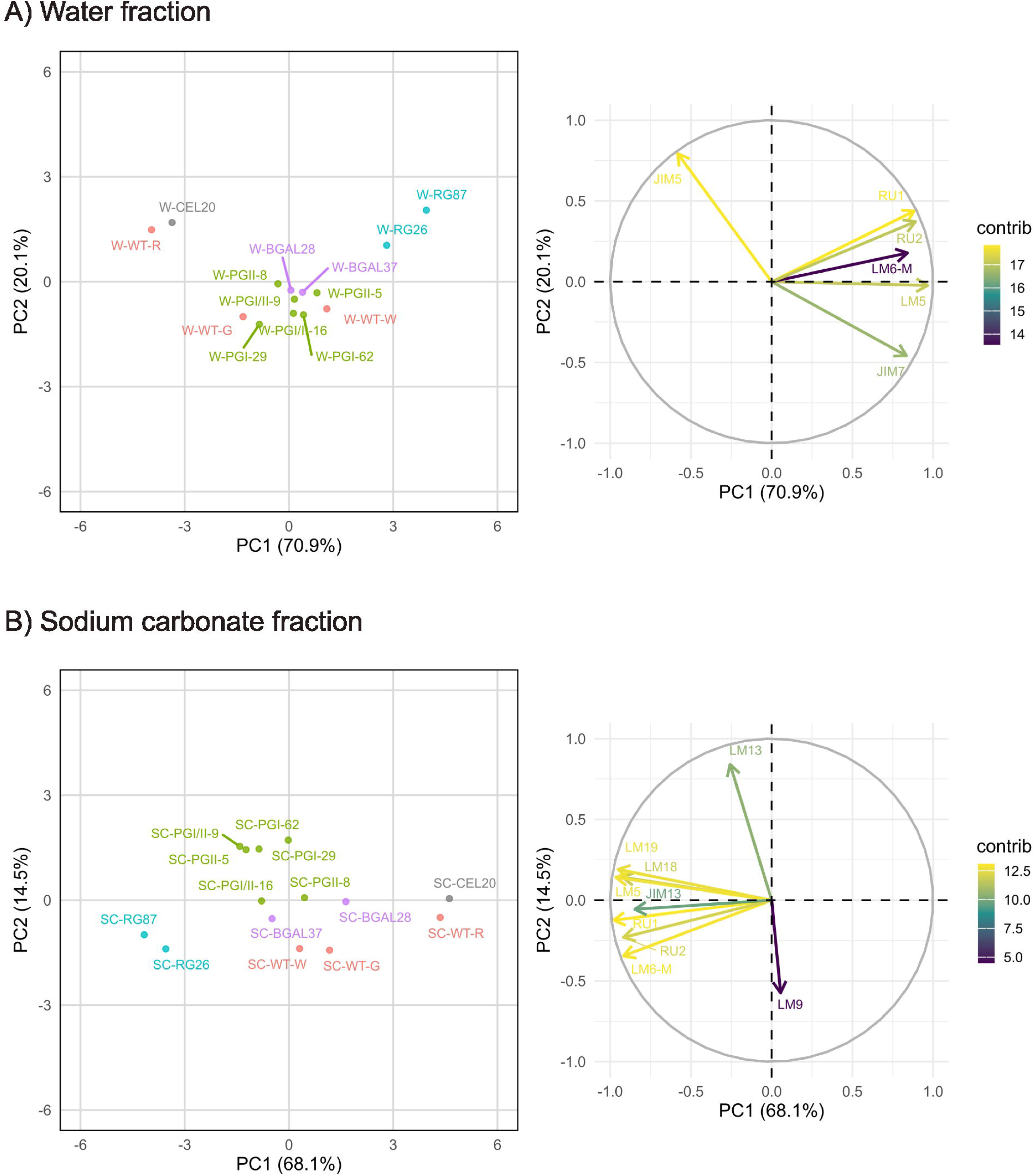
PCAs of the carbohydrate microarray data derived from the water (A) and sodium carbonate (B) cell wall fractions. The factor score plots appear on the left, whereas the variable plots are shown on the right. The color scale in the variable plots reflects the average contribution of each variable to the variation accounted for by the two principal components.

**Figure 7.**
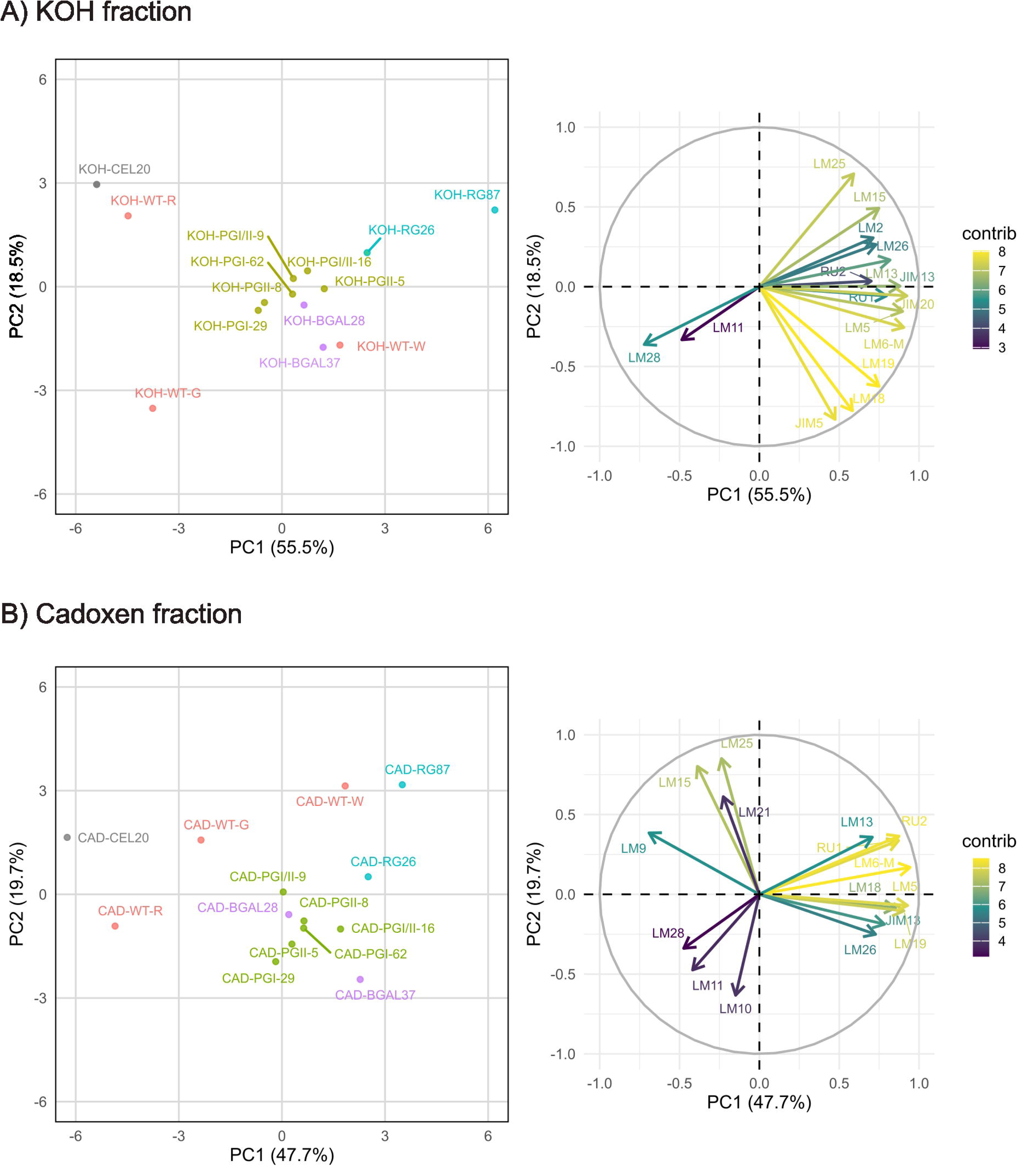
PCAs of the carbohydrate microarray data derived from the KOH (A) and cadoxen (B) cell wall fractions. The factor score plots appear on the left, whereas the variable plots are shown on the right. The color scale in the variable plots reflects the average contribution of each variable to the variation accounted for by the two principal components.

Correlation matrices of relative absorbance values for mAbs in each fraction are presented in Figure 8. In the water fraction, methyl-esterified HG (JIM7) correlated negatively with low methyl-esterified HG (JIM5), while RG-I-related mAbs (RU1, RU2, LM5, LM6-M) were all positively correlated among them and with methyl-esterified HG. HG and RG-I mAbs were positively correlated in the remaining fractions. Pectin epitopes, especially branched galactans (LM26), were positively correlated with AGP glycan epitopes (JIM13). In the KOH fraction, xyloglucans (LM15, LM25) correlated positively with galactans (LM5, LM26), extensins (JIM20), and AGPs (JIM13), and negatively with xylans (LM11, LM28). Heteroxylan epitopes (LM11, LM28) were negatively correlated with RG-I (RU, LM6, LM13) in both KOH and cadoxen fractions. Finally, the average increase in fruit firmness of ripe fruits among transgenic lines was incorporated into the correlation analysis. The increase in firmness correlated positively with a higher proportion of partially methyl-esterified HG in the water fraction (positive correlation coefficient with JIM7 abundance and negative correlation coefficient with JIM5), a higher abundance of HG pectins (JIM5, LM18 and LM19) in the rest of fractions and RG-I galactan epitopes (LM5) in the cadoxen fraction, and a lower abundance of galactosylated xyloglucans (LM25) in the cadoxen fraction.

**Figure 8.**
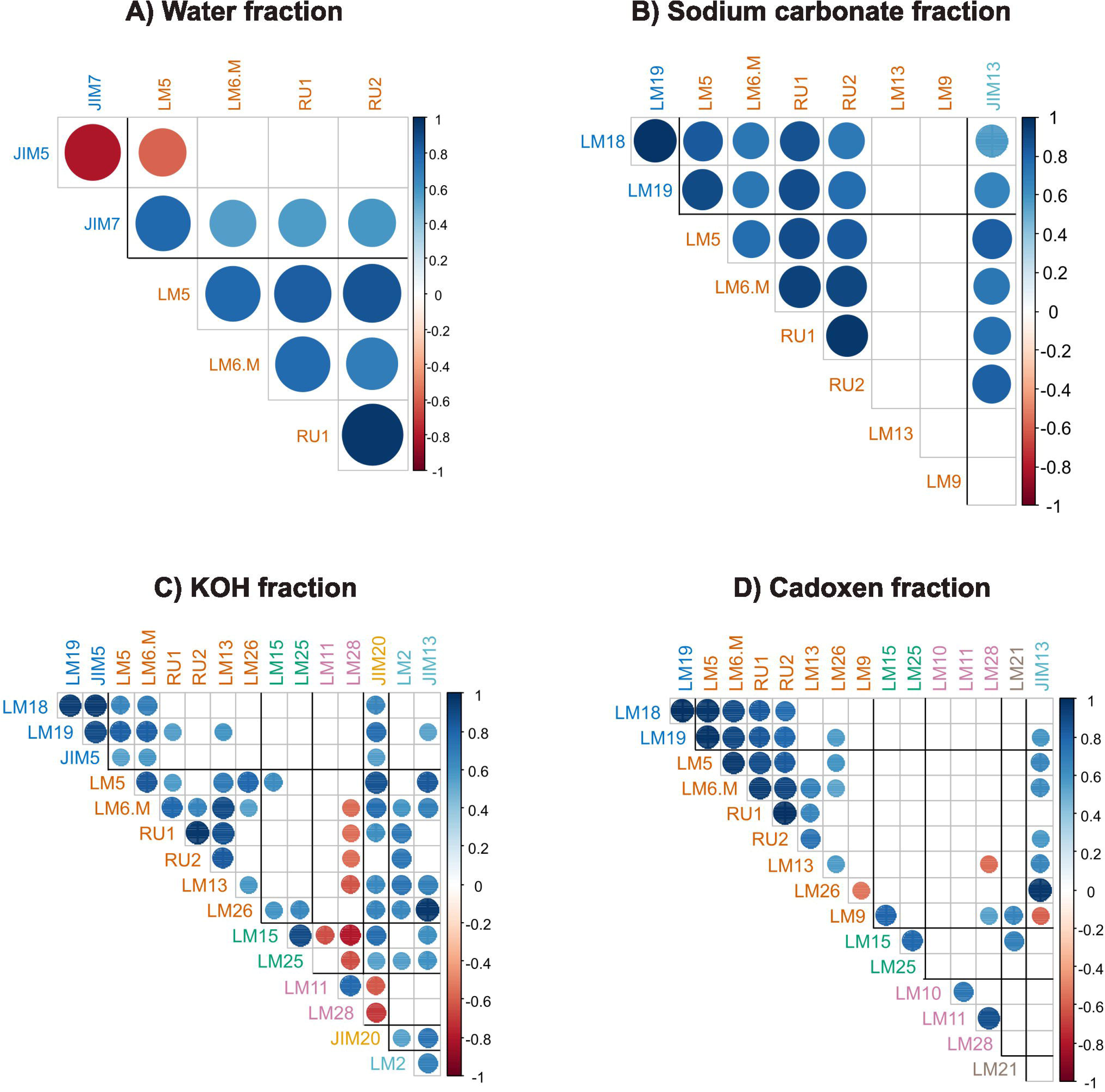
Correlation plots of mAb relative abundance across the different cell wall fractions of ripe fruits. Only significant correlations (P=0.05) are displayed.

## 4. Discussion

### 4.1. The firmer fruit phenotype is stably maintained through vegetative propagation

The meta-analysis of fruit quality traits in transgenic and control strawberry plants across different growing seasons revealed that variability in fruit firmness among pectinase mutants and control lines was primarily explained by genotype and harvest year. In contrast, fruit morphological traits (weight, length, and width) and soluble solids content (SSC) had negligible effects. The absence of significant correlations between firmness and fruit size or phenotypical parameters indicates that the increase in firmness observed in transgenic lines is not a secondary consequence of altered fruit development or sugar accumulation, but rather a direct outcome of modifications in cell wall metabolism due to down-regulation of pectin-degrading enzymes. The fitted model, which explains nearly half of the variability in firmness (adjusted R² = 0.481), identified significant positive coefficients for all transgenic genotypes compared to the wild type. However, the low number of fruits produced by some of the transgenic lines in certain harvest years may have influenced the results obtained. This confirms that silencing of pectinase genes consistently enhances firmness, in agreement with previous reports. The genotype-dependent variation observed here, where antisense PG lines showed the most potent effects, B-Gal an intermediate behaviour, and RGLyase lines displayed the lowest, likely reflects the potential impact of these enzymes and their cell wall targets in the softening process. Enzyme redundancy among pectinase isoforms could also explain these differences in the firmer phenotype.

On the other hand, the significant effect of harvest year, with 2013 yielding the firmest fruit and 2021 the softest, could point to environmental modulation of firmness, possibly through temperature. The addition of different temperature indexes, such as the average temperature at harvest of each fruit or the average temperatures of the 15 days preceding harvest, had a lower effect on the model. Although these temperature indexes displayed a low negative correlation with firmness, the adjusted R² of the model was not improved. In agreement with these results, Hopf et al. (2022) correlated different weather indices with strawberry fruit quality. They found that preharvest temperature decreased the content of soluble solids and titratable acidity but had no effect on fruit firmness. Then, other uncontrolled factors, such as annual variations in plant management, including pest control treatments, plant physiological status, or the operator’s measurement of fruit quality variables, could account for the variability observed over the years. Undesired transgene silencing is of significant concern when using transgenic approaches for crop improvement (Rajeevkumar et al., 2015). Along this line, few studies have analysed the stability of a transgenic phenotype over different years, particularly in clonally propagated plants. The persistence of the firmer phenotype across all transgenic genotypes over multiple years indicates that the pectinase suppression phenotype is stable and robust to environmental variation and clonal propagation of strawberry plants.

### 4.2. Down-regulation of pectinase genes impairs cell wall disassembly

The silencing of PGs, β-Gal, and RGLyase genes significantly altered the cell wall carbohydrate profile of ripe fruits when compared with the control. These changes were similar in the three kinds of pectinase mutants but subtle differences in fruit texture were observed among these lines, as previously described. In general, the microarray profile of ripe fruits from PG and RGLyase lines were similar to those previously described (Paniagua et al., 2020; Ric-Varas et al., 2024). By contrast, endo-β-glucanase silencing did not modify the cell wall disassembly process and the microarray profile was quite like control fruits.

Polygalacturonases catalyze the hydrolytic cleavage of galacturonic linkages in HG domains between demethylated galacturonic acid residues. As expected, there were minor differences in the microarray signals of PG mutants, including both single *FaPG1* and *FaPG2* mutants, and double *FaPG1*-*FaPG2* mutants. Interestingly, down-regulation of PG genes not only increased the amount of HG epitopes detected with JIM7, LM18 and LM19, but also the abundance of RG-I backbone (RU1 and RU2 mAbs) and side chain epitopes (LM5 and LM6 mAbs). Although the chemical composition of pectins is well known, the way in which pectin domains are interconnected within the cell wall is unclear. The most widely accepted model depicts the pectin composite as a linear backbone composed of HG (smooth regions), which can be interspersed with rhamnosyl residues, alternating at regular intervals with branched RG-I, xylogalacturonan (XGA), arabinan, and arabinogalactan (hairy regions) (De Vries et al., 1982). Alternative models postulate RG-I as the main pectin backbone decorated with XGA and linear HG as lateral (Vincken et al., 2003) or terminal (Yapo, 2011) chains. The analysis of strawberry fruit pectins by AFM suggested the presence of a mixture of HG chains with a reduced number of branches and complex structures supposedly formed by an HG unit linked to an RG-I core (Paniagua et al., 2017b). That result is in accordance with the De Vries pectin model and could explain the increased abundance of RG-I epitopes in PG mutants found in this research. On the other hand, Oechslin et al. (2003) identified a pectin domain in apple fruit associated with cellulose, containing an RG-I backbone with XGA, a linear galactan, and highly ramified arabinan side chains. HG cannot bind to cellulose in vitro, while galactan and arabinan showed moderate binding capacity, although substantially lower than xyloglucans (Zykwinska et al., 2005, 2008); therefore, the presence of HG in KOH and cadoxen fractions from antisense PG strawberry fruits could be explained if these pectin domains were linked to xyloglucan or cellulose through RG-I. Notably, Cornuault et al. (2018) detected LM5 galactan and RU1 backbone epitopes in the KOH fractions of four kinds of fruits, being higher in those with a firmer texture, aubergine and apple, than in soft fruits, tomato and strawberry.

β-galactosidases hydrolize terminal non-reducing β-D-galactosyl residues from β-D-galactoside substrates (Tateishi, 2008). A common feature of the cell wall remodeling during fruit ripening is the loss of galactose from pectin side chains, supposedly due to the activation of β-galactosidases (Brumell, 2006). The sugar composition analysis of ripe antisense *Fa*β*Gal4* lines revealed a 30% increase in Gal content in the transgenic lines, with the KOH cell wall fraction displaying the highest increment in Gal (Paniagua et al., 2016). The results obtained in the microarray support these previous findings. The abundance of the LM5 epitope, specific against β-D-galactan (Jones et al., 1997), was higher in βGal transgenic lines in all cell wall fractions when compared with the control fruit at unripe or ripe stages. However, the highest signals of this mAb were found in RGLyase mutants. The breakdown of RG-I backbone by RGLyases could facilitate the access of β-galactosidases to their substrate, contributing to Gal release during softening. Globally, the microarray profile of BGal lines did not differ significantly from that of PG mutants except for a slightly lower abundance of HG epitopes in the SC fraction. This result reinforces the link between HG and RG-I *in muro* previously discussed.

RGLyases are the less studied pectinase family, and functional analysis of *RGLyase* genes in fleshy fruits is scarce. These enzymes potentially break the backbone of RG-I (McDonough et al., 2004). Its silencing in RGLyase mutants significantly enhanced the amount of RG-I epitopes, both RG-I backbone (RU1 and RU2 mAbs) and galactan and arabinan side chains (LM5 and LM6 mAbs, respectively). This increase was evident in all fractions, particularly in pectin-enriched fractions (water and SC) and KOH. Notably, these mutants displayed a decreased amount of xylan epitopes (LM28) but an increased amount of AGP (JIM13).

The factor score plots of the PCA analysis provide a global picture of the changes induced by pectinase gene silencing in strawberry fruit. In all cell wall fractions, the mutant plants appeared close to control fruits at unripe developmental stages. The expression of these target genes was induced by ABA and reached its peak at the white-red stage (Moya-León et al., 2019). Clearly, their silencing impaired cell wall remodeling, and it had an impact on fruit firmness. RGLyase fruits displayed the most different microarray profile among all cell wall mutants, but the lower increase in fruit firmness. It can be suggested that sub-populations of HG and RG-I pectins are tightly linked to the cell wall, contributing to the mechanical properties of strawberry fruit at unripe stages; the removal of these pectins could contribute to cell wall weakening and strawberry softening.

The main changes in the cell wall of the evaluated pectinase mutants can be summarized as follows: (i) a higher proportion of methylated to non-methylated HG in the water-soluble fraction; (ii) an increase in the abundance of RG-I across all cell wall fractions, including those enriched in xyloglucan (KOH) and in matrix glycans associated with cellulose (cadoxen); and (iii) an increase in AGP epitopes, particularly in the xyloglucan fraction. The carbohydrate microarray revealed that HG pectins from the water-soluble fraction underwent a gradual demethylation process from the green to the red stages, consistent with the induction of specific pectin methylesterase (PME) genes during fruit ripening (Castillejo et al., 2004; Xue et al., 2020). Down-regulation of PG, β-Gal, or RGLyase genes impaired this demethylation process. RNA-seq analyses of ripe fruits from both *FaPG* and *RGLyase* mutants showed that the expression of other genes encoding cell wall–modifying enzymes was also altered. The reduced HG demethylation may be associated with the down-regulation of pectinesterase genes, as reported in the RNA-seq study of *FaPG* mutants (Paniagua et al., 2020), or with the silencing of genes encoding proteins involved in cell wall integrity and remodeling, such as xyloglucan endotransglucosylase/hydrolase (XTH), as observed in *RGLyase* plants (Ric-Varas et al., 2024). A comparative transcriptomic analysis of pectinase mutants would help clarify this aspect. On the other hand, AGPs have been implicated in a wide range of physiological processes, including germination, cell expansion, cell signaling, modulation of cell wall mechanics, and pectin assembly (Hijazi et al., 2014; Leszczuk et al., 2018). In apple fruit, AGP epitopes detected with JIM13 were localized in the inner cell wall in association with the plasma membrane. Their abundance increased from the green to the ripe stages, but later declined during storage, concomitant with the disruption of the cell wall-plasma membrane continuum (Leszczuk et al., 2018). An increase in AGP epitopes has also been reported during grape ripening (Moore et al., 2014). In strawberry, the JIM13 epitope likewise increased during ripening, but its abundance was significantly higher in the transgenic pectinase mutants. This may result from reduced pectin degradation, as it has been proposed that AGPs can be covalently linked to RG-I and HG within a proteoglycan complex, potentially forming a continuous network between wall polysaccharides and structural wall proteins (Tan et al., 2013; Mohnen et al., 2023). Further research is needed to elucidate the role of this complex in fruit softening.

## 5. Conclusions

The results of this study demonstrate that suppressing pectinase genes in strawberry leads to a stable enhancement of fruit firmness, independent of harvest year. Multiple linear regression analysis confirmed that genotype and harvest year were the main factors influencing firmness, while fruit size and soluble solids content played no significant role. All transgenic lines exhibited a consistently firmer phenotype compared with wild-type fruits, underscoring the robustness of the trait over time. On the other hand, cell wall compositional analyses revealed that the firmer phenotype is associated with reduced de-methylesterification of HG and the retention of RG-I backbone and side-chain epitopes in ripe fruits. These characteristics indicate a partial inhibition of cell wall disassembly during ripening and the structural support role of pectins in the wall. Principal component and correlation analyses further supported a close relationship between the accumulation of specific pectic epitopes and the maintenance of fruit firmness. Overall, our findings provide clear evidence that targeted suppression of pectinase activity modulates pectin remodeling pathways, resulting in a durable modification of cell wall structure and texture properties. This work advances our understanding of the molecular determinants of fruit softening and provides a promising strategy for enhancing postharvest texture in strawberries and potentially other fleshy fruits.

## Supplementary Material

**Supplementary Figure 1.** Aspect of plants and ripe fruits from some pectinase lines

**Supplementary Figure 2.** Bivariant plots of fruit variables.

**Supplementary Figure 3.** Diagnosis of the multiple linear regression models

**Supplementary Table 1.** Number of fruits evaluated per genotype and year

**Supplementary Table 2.** List of monoclonal antibodies used in the carbohydrate microarray

