## Supplementary material for "Suppression of Pectinase Genes Confers Stable Enhancement of Fruit Firmness through Modulation of Pectin Structure in Strawberry": Supplementary_Material.pdf

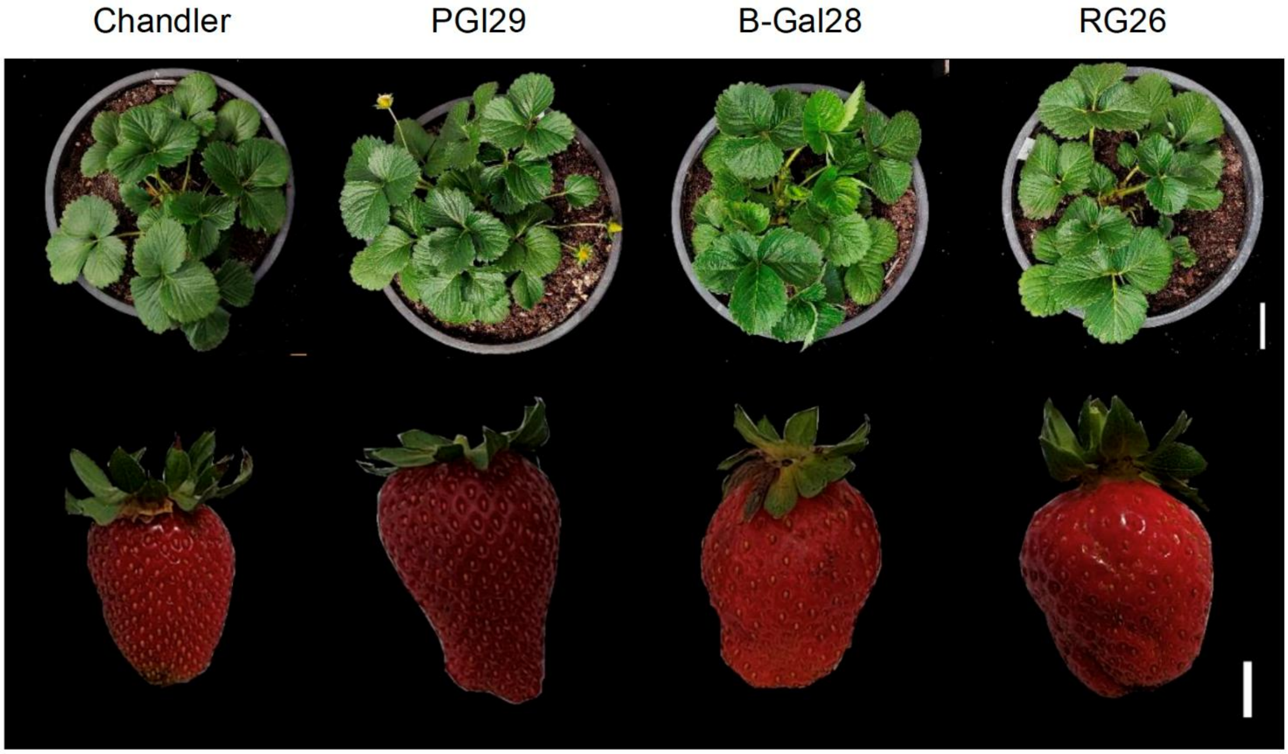


**Supplementary Figure 1.** Aspect of plants and ripe fruits from control (Chandler) and some selected pectinase lines. Bars correspond to 5 cm (upper bar) and 1 cm (lower bar).

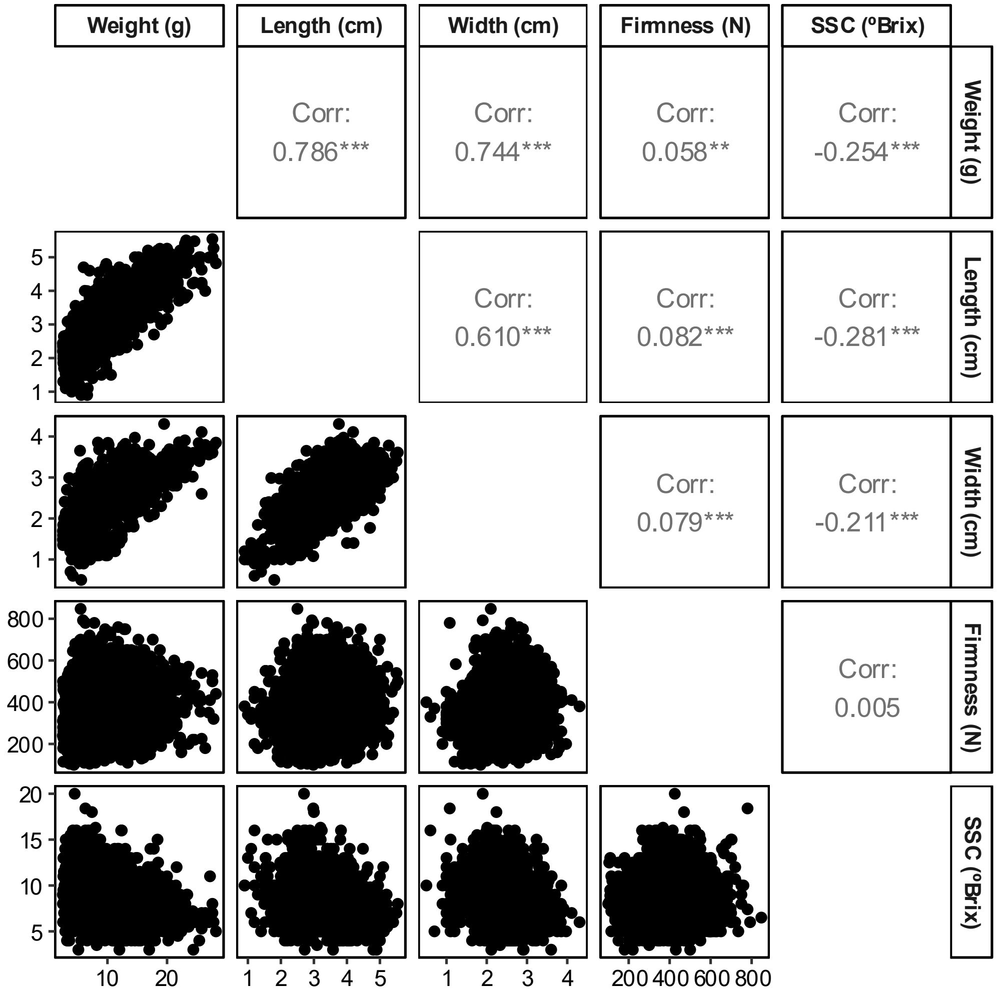


**Supplementary Figure 2.** Bivariant plots showing the relationship between fruit weight, length, width, firmness and soluble solid content (n = 2856). Correlation (corr.) between variable pairs was evaluated using Pearson's linear coefficient that was contrasted using a *t*-test. Asterisks denote a significant difference: * P-value < 0.05, ** P-value < 0.01, and *** P-value < 0.001.

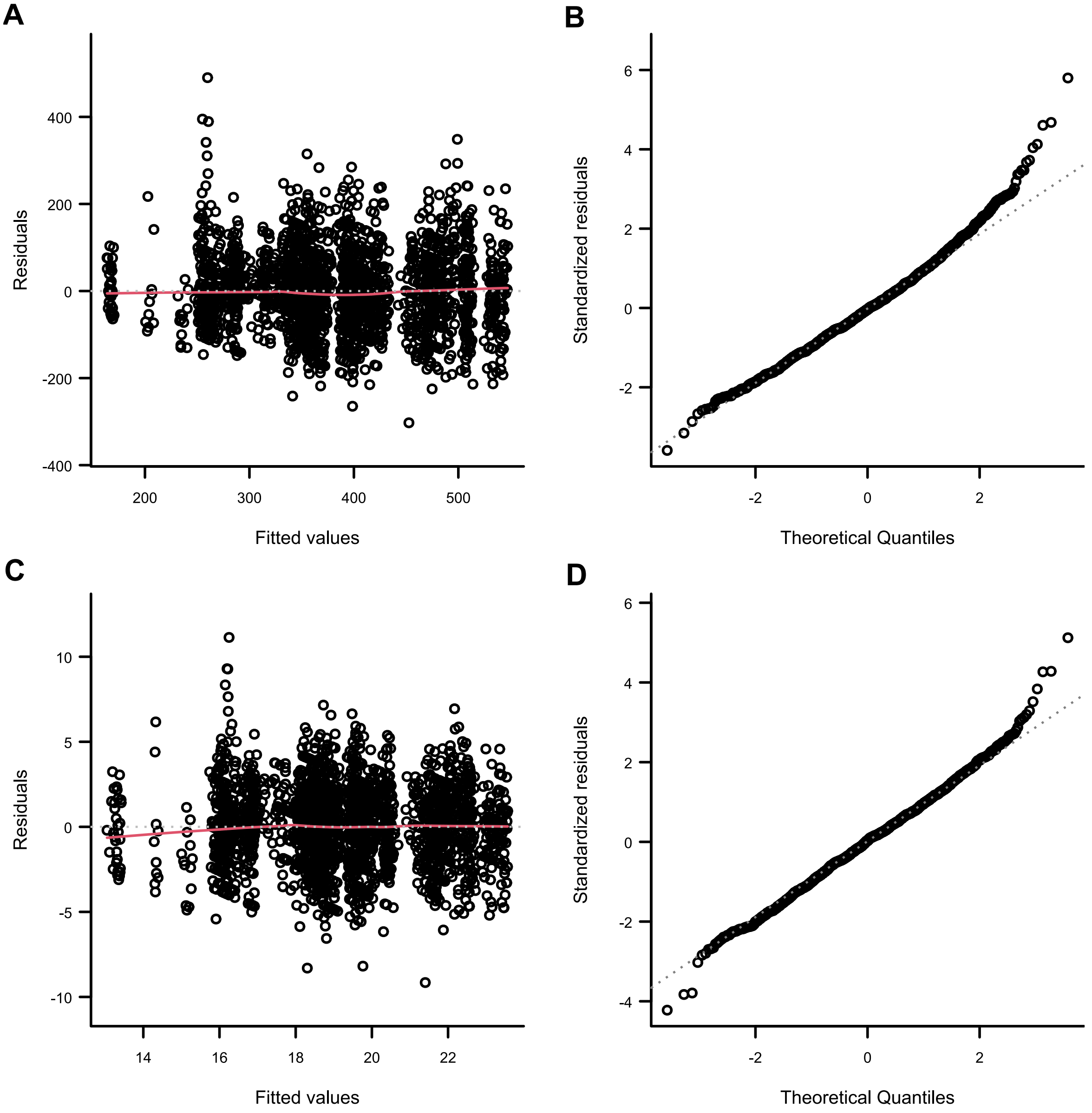


**Supplementary Figure 3.** Diagnosis of the initial (A, B) and final (C, D) multiple linear regression models. Initial model was adjusted using fruit firmness as response variable and fruit length, width, soluble solid content, harvest year and genotype as response variables. In the final model, root square of fruit firmness was used as response variable and all numerical variables were standardized. A and C show the residuals vs the fitted values of the models. B and D show the Q-Q plot of the residuals of the models.

**Supplementary Table 1.** Number of fruits evaluated per genotype and year. NA, transgenic genotypes not available that year. -- Not included in the analysis due to the low number of fruits obtained that year

| | Genotype | | | | | | | | | | |
|---|---|---|---|---|---|---|---|---|---|---|---|
| **Year** | **WT** | **PGI-29** | **PGI-62** | **PGI/II-9** | **PGI/II-16** | **PGII-5** | **PGII-8** | **RG26** | **RG87** | **β-Gal28** | **β-Gal37** |
| **2011** | 261 | 103 | 85 | NA | NA | NA | NA | NA | NA | 28 | 32 |
| **2012** | 151 | 36 | -- | 25 | 11 | 20 | 13 | NA | NA | 51 | 60 |
| **2013** | 147 | 65 | -- | 61 | 63 | 48 | 38 | NA | NA | 61 | 51 |
| **2014** | 82 | 78 | -- | 62 | 60 | 37 | 52 | NA | NA | -- | -- |
| **2015** | 30 | 11 | -- | -- | -- | -- | 8 | 18 | 11 | -- | -- |
| **2016** | 214 | 48 | 20 | 26 | 21 | 27 | -- | 31 | 38 | | -- |
| **2021** | 33 | 12 | 13 | -- | 13 | -- | 9 | 13 | 10 | -- | -- |
| **2023** | 52 | 70 | -- | -- | -- | -- | -- | 68 | -- | 82 | -- |
| **2024** | 115 | 41 | -- | -- | -- | -- | -- | 20 | -- | 21 | -- |

**Supplementary Table 2.** List of monoclonal antibodies used in the carbohydrate microarray and the epitopes recognized by each mAb.

| **Antibody** | **Cell wall epitope** | **Reference** |
|---|---|---|
| **Homogalacturonan (HG)-related** | | |
| LM18 | Partially Me-HG / no ester | Verhertbruggen et al. (2009) Carbohyd. Res. 344, 1858 |
| LM19 | Partially Me-HG / no ester | Verhertbruggen et al. (2009) Carbohyd. Res. 344, 1858 |
| JIM5 | Partially Me-HG / no ester | VandenBosch et al. (1989) EMBO Journal 8, 335-342 |
| LM7 | Partially Me-HG / non-blockwise | Willats et al. (2001) J. Biol. Chem. 276, 19404-19413 |
| JIM7 | Partially Me-HG | Knox et al. (1990) Planta 181, 512-521 |
| LM20 | partially Me-HG | Verhertbruggen et al. (2009) Carbohydr. Res. 344, 1858 |
| LM8 | xylogalacturonan | Willats et al. (2004) Planta 218, 673-681 |
| **Rhamnogalacturonan I (RGI)-related** | | |
| LM5 | (1→4)-β-D-galactan | Jones et al. (1997) Plant Physiol. 113, 1405-1412 |
| LM6 | (1→5)-α-L-arabinan | Cornuault et al. (2017) BiorXiv: doi.org/10.1101/161604 |
| RU1 | [→2)-α-L-rhamnose- (1→4)-α-D-galacturonic acid-(1→]$_7$ | Ralet et al. (2010) Planta, 231, 1373-1383 |
| RU2 | [→2)-α-L-rhamnose- (1→4)-α-D-galacturonic acid-(1→]$_7$ | Ralet et al. (2010) Planta, 231, 1373-1383 |
| LM13 | Linearised (1→5)-α-L-arabinan | Moller et al. (2008) Glycoconjugate J. 25, 37-48 |
| LM26 | Branched galactan | Torode et al. (2018) Plant Physiol. 176, 1547-1558 |
| LM9 | feruloylated (1→4)-β-D-galactan | Clausen et al. (2004) Planta 219, 1036-1041 |
| LM16 | processed arabinan/put. galactan stub | Verhertbruggen et al. (2009) Plant Journal 59, 413-425 |
| **Xyloglucan** | | |
| LM15 | Xyloglucan (XXXG motif) | Marcus et al. (2008) BMC Plant Biol. 8, 60 |
| LM24 | Galactosylated xyloglucan | Pedersen et al. (2012) J. Biol. Chem. 287, 39429-39438 |
| LM25 | XXXG/galactosylated xyloglucan | Pedersen et al. (2012) J. Biol. Chem. 287, 39429-39438 |
| **Heteroxylan** | | |
| LM10 | (1→4)-β-D-xylan | McCartney et al. (2005) J. Histochem. Cytochem. 53, 543 |
| LM11 | (1→4)-β-D-xylan /arabinoxylan | McCartney et al. (2005) J. Histochem. Cytochem. 53, 543 |
| LM28 | Glucuronoxylan | Cornuault et al. (2015) Planta 242, 1321-1334 |
| LM27 | unknown epitope assoc. grass xylan | Cornuault et al. (2015) Planta 10.1007/s00425-015-2375-4 |
| LM12 | ferulic acid, feruloylated xylan | Pedersen et al. (2012) J. Biol. Chem. 287, 39429-39438 |
| **Heteromannan** | | |
| LM21 | Heteromannan | Marcus et al. (2010) Plant J. 64, 191-203 |
| **Extensin** | | |
| JIM20 | Extensin | Smallwood et al. (1994) Plant J. 5, 237-246 |
| JIM11 | extensin | Smallwood et al. (1994) Plant Journal 5, 237-246 |
| **Arabinogalactan-protein glycan (AGP)** | | |
| JIM13 | AGP glycan | Knox, et al. (1991) Plant J. 1, 317-326 |
| LM2 | β-linked-GlcA in AGP glycan | Yates et al. (1996) Glycobiology 6, 131-139 |
| JIM14 | AGP glycan | Knox, et al. (1991) Plant Journal 1, 317-326 |
| JIM15 | AGP glycan | Knox, et al. (1991) Plant Journal 1, 317-326 |
| JIM16 | AGP glycan | Knox, et al. (1991) Plant Journal 1, 317-326 |
| **Other cell wall Abs** | | |
| LM23 | non-acetylated xylosyl in xylogalacturonan, xylan, fucoidan preps | Manabe et al. (2011) Plant Physiology 155, 1068-1078 |
